# Exploring the genetic factors of nitrogen use efficiency in potato Genetics of Nitrogen Use Efficiency in Potato

**DOI:** 10.1101/2025.05.18.654744

**Authors:** Miguel Ángel Mendoza-Bustamante, Aura Natalia Jiménez-Medrano, Johana Carolina Soto-Sedano, María Cecilia Delgado-Niño, Stanislav Magnitskiy, Gustavo Adolfo Ligarreto-Moreno, Teresa Mosquera-Vásquez

## Abstract

Nitrogen is an essential nutrient for plants, used by farmers to increase the yield of the crops. However, this practice increases greenhouse gases, negatively affecting the environment. Nitrogen Use Efficiency (NUE) is a trait that is beginning to be studied in some model species and in cereals due to its complex and novel trait nature. In potatoes, the information is scarcer. The study of NUE at the genetic level, based on a diverse population in potato materials, will contribute to the understanding of the genetic architecture of the trait. This research evaluated NUE in a *Solanum tuberosum* diploid potato genetic diversity panel from the Phureja group. The characterization of the trait was carried out in substrate conditions, for low and high levels of nitrogen for both the vegetative and the reproductive phase. Eighteen variables associated with NUE were measured, nine under low nitrogen and nine under high nitrogen conditions. A GWAS was conducted, and a total of 21 QTNs were identified as strongly associated with 11 phenotypic variables related to NUE in potato, along with a repertoire of 750 candidate genes associated with the trait. This research aimed to establish the basis for understanding the genetic architecture of NUE in *Solanum tuberosum*. Likewise, the knowledge derived is expected to be useful for plant breeding programs that begin to improve this trait.

## Introduction

Potato is an essential food for humanity, characterized by its high nutritional value, significant yield potential, and wide adaptability, making it a crucial crop for global food security (Koch *et al.,* 2020). It is considered an important source of carbohydrates, but it also provides dietary fiber, vitamin C, vitamin B6, potassium, magnesium, iron, carotenoids, and phenolic acids (Hu, *et al.,* 2024, Riveros-Loaiza *et al.,* 2022; Parra-Galindo *et al.,* 2019; Narváez-Cuenca *et al.,* 2018). It is affordable food that is accessible throughout the year. The dominant approach in potato production has followed a linear model, assuming that higher inputs lead to higher yields. This model has resulted in the overuse of fertilizers, especially nitrogen (N), which has lowered efficiency and contributed to water pollution and soil degradation (Su *et al.,* 2024). This approach mirrors the broader trends seen since the Green Revolution, which, following the end of World War II, introduced synthetic chemical pesticides and fertilizers. These products led to increased crop yields, with wheat yields increasing by up to 400% (Chilon, 2017; Nasholm *et al.,* 2009). This positioned nitrogen as one of the most important fertilizers in crop production. In the last 20 years, there has not been a proportional increase in crop yields despite increased applications of nitrogen fertilizers (Sheng *et al.,* 2013). It is reported that between 50% to 75% of the nitrogen applied as fertilizer is not utilized by plants, instead leaching as nitrates *(NO* ^—^) or volatilizing as ammonia *(NH_3_)* and nitrous oxide (*N_2_O)*, thus polluting the environment (Hirel and Krapp, 2020; Li *et al.,* 2022a). As a result, Nitrogen Use Efficiency (NUE) has become the second most studied trait worldwide due to its importance in agriculture for optimizing fertilization and its potential for breeding programs (Getahun *et al.,* 2020). However, the genetic control of this trait remains largely unexplored, as NUE is a complex trait governed by multiple genes involved in control. To reduce the use of nitrogen fertilizers in crop production, efforts have been made to decipher the architecture of the traits in crops such as maize, wheat, and rice (Siddiqui *et al.,* 2023; Zang *et al.,* 2024; Yu *et al.,* 2025). Genome-wide association studies (GWAS) are a valuable strategy for identifying genomic regions associated with a trait of interest. Using a naturally diverse population enhances the likelihood of identifying superior alleles in the gene pool and provides more precise mapping (Sood *et al.,* 2024). Genetic association studies in NUE have identified candidate genes such as *ZmNAC36* for maize (Wang *et al.,* 2022), *NPF2.12* for wheat (Siddiqui *et al.,* 2023), and *OsNIGT1* for rice (Li *et al.,* 2022), among others. Understanding the architecture of the traits in rice has improved NUE by up to 40% with the overexpression of the *OsNRT2.3b* gene, enhancing the buffering capacity of the plant cell, to increase nitrogen uptake (Fan *et al.,* 2017).

In the case of potato, NUE studies have mainly focused on improving cultural practices or soil management, except for some researchers who have conducted genetic studies using linkage mapping (Getahun *et al.,* 2020) and association mapping (Getahun *et al.,* 2022; Ospina *et al.,* 2021). Although these studies have found QTL associated with NUE, candidate genes controlling the trait have not been identified. This may be due to the methodologies used for phenotyping, as studies were conducted under field conditions where precipitation, temperature, and their interaction with the soil, rich in minerals and microorganisms can affect nitrogen availability (Geng *et al.,* 2018; Ma *et al.,* 2012). These conditions affect the proposed treatments. It may also be due to insufficient genetic diversity in the studies to reveal variants associated with NUE, as those studies have been conducted in biparental populations contrasting in precocity and in commercially European populations. Therefore, it is necessary to conduct studies with highly diverse populations that allow the study of natural allelic diversity, with phenotyping in controlled environments that establish conditions of nitrogen deficiency and sufficiency, to help decipher the genetic architecture of the trait, generating knowledge about the genetic architecture of NUE in a diversity panel of diploid potato through genetic association analysis. In this context, this study aimed to identify genetic variants associated with NUE in a potato population using GWAS, with the goal of contributing to genetic improvements in potato for optimized NUE. To achieve this, NUE was evaluated in a diploid population from the Phureja group, selected for its genetic and nutritional potential (Juyó *et al.,* 2015; Parra Galindo *et al.,* 2021), making it an excellent candidate for investigating genetic variability and enhancing this trait. The QTNs (Quantitative Trait Nucleotides) and candidate genes identified potentially associated with NUE, enhancing our fundamental understanding of the genetic mechanisms underlying the NUE, which could guide future genetic breeding strategies to optimize this trait in potatoes.

## Materials and methods

### Plant material and essay conditions

A total of 100 accessions (genotypes) of diploid potato, *Solanum tuberosum* Phureja group, from the Working Collection of the Potato Breeding Program of the Universidad Nacional de Colombia, plus five diploid commercial varieties: Criolla Colombia, Criolla Galeras, Criolla Guaneña, Criolla Latina, Criolla Paisa, were used as association panel (S1 Table). The essay was established in the greenhouses of the Faculty of Agricultural Sciences of the Universidad Nacional de Colombia (4°38’16.7 “N, 74°05’18.2” W). During the crop cycle the average air temperature was 19.3 °C, with a maximum and the minimum temperature of 38.8°C and 8°C respectively. The average relative air humidity was 63.6% and the average dew point was 11.38°C.

The essay was carried out under a completely randomized design (CRD), with 2 treatments and 3 biological replicates (plants) per genotype and treatment. The plants were grown in plastic bags with a 4 kg substrate capacity. The substrate used was a mixture (v/v) of 70% of peat (Klasmann^®^ No. 413 peat, 0-5 mm, without added nutrients) and 30% perlite (8-12 mm). This system, as a Semi-aeroponic system, allows for controlled nutrient supply without affecting tuber development.

### Phenotyping Nitrogen Use Efficiency (NUE) parameters in potato

#### Establishment of nitrogen deficient and sufficient treatments

Treatments (T) consisted of a low nitrogen dose (nitrogen-deficient conditions) and a sufficient nitrogen dose (high nitrogen conditions), applied at 2 crop stages: vegetative and reproductive stage. For the vegetative stage, the dose of low nitrogen was 19 ppm and 195 ppm for high nitrogen, while, for the reproductive stage, there was low nitrogen 15 ppm and high nitrogen 146 ppm. Nitrogen was supplied as ammonium nitrate (NH_4_NO_3_) of analytical grade purity. Total nitrogen supply per plant per growth cycle, including nitrogen content in water and substrate, was 0.56 g (low nitrogen) and 1.78 g (high nitrogen).

The dosages of each macronutrient and micronutrient (in ppm), excluding nitrogen, were established for the vegetative and reproductive stages following the criteria reported by Jiménez-Medrano *et al*. (*manuscript under review*): P (35, 22), K (200, 260), Ca (64, 64), S (40, 40), Mg (24, 24), B: (0.6, 0.6), Zn (0.2, 0.2), Cu (0.2, 0.2), Mn (0.75, 0.75), Fe (1.5, 1.5), Mo (0.04, 0.04). The nutrient solution was applied twice a week via the irrigation water, adjusting to the water needs of the plants and substrate moisture. In addition, the electrical conductivity of the substrate was monitored and maintained at approximately 1.5 dS⋅cm^−1^. The volume of solutions applied to each plant was 8.6 L in total, 2.4 L in the vegetative stage and 6.4 L in the reproductive stage. The transition from the vegetative to the reproductive stage nutrient solution occurred 49 days after planting (DAP), corresponding to 556.1 growing degree days (GDD) calculated using the equation from Arnold (1959).

#### Measurement of variables linked to nitrogen use efficiency

A total of 18 phenotypic variables were measured per genotype across 3 replicates, 9 under high N treatment and 9 under low N treatment. At 110 DAP, which corresponded to the time of harvest, the following variables were measured in each plant: 1) Relative chlorophyll content (SPAD74) at 74 days after planting (DAP) as an indicator of the N status of the plants with a portable Minolta SPAD 502^®^ (Japan). Readings were taken in leaves as the median of 3 fully expanded leaves, in the upper third of each evaluated plant; 2) Number of stems (StemNo). 3) Stem length (StemL), was measured as the length of the longest stem on each plant, from the base where it joins the root system to the shoot apex; 4) Number of tubers (TubNo).

Samples of the aerial organs and tubers were collected separately and dried in an oven at 60 °C for 3 to 5 days until a constant weight was reached. The dry weight of the aerial part and the tubers was determined individually, and their sum was used to calculate the total dry weight. Also, samples of aerial organs (leaves and stems) were analyzed for Carbon percentage of the aerial part of the plant (%CAp) and tubers were analyzed for total nitrogen percentage (%Ntub), both using DUMAS methodology with the Leco Fp828 mc 50282^®^ Series Elemental Analyzer. These analyses were conducted at the Water and Soil Laboratory of the Faculty of Agricultural Sciences, Universidad Nacional de Colombia, Bogotá. Finally, the NUE indices described by Zebarth *et al*. (2008) and Tiwari *et al*. (2020) were calculated as follows: Nitrogen Use Efficiency (NUE, g⋅g^−1^): dry weight of tuber/N supplied. Agronomic Nitrogen Use Efficiency (AgNUE, g⋅g^−1^): Fresh weight of tubers/N supplied. Harvest Index (HI, g⋅g^−1^): dry weight of tubers/dry weight of total plant.

Phenotypic data were subjected to analysis of variance (ANOVA) within and between genotypes, followed by the Tukey’s test. To explore multivariate patterns within the phenotypic dataset and identify traits most strongly associated with NUE), a Principal Component Analysis (PCA) was performed using scaled phenotypic data collected from potato genotypes evaluated under contrasting nitrogen supply conditions. The analysis was conducted in R, using the FactoMineR (Lê *et al.,* 2008) and factoextra (Kassambara *et al.,* 2016) packages. PCA enabled dimensionality reduction and visualization of genotype distribution across multiple traits related to NUE performance.

To identify genotypes with contrasting Agronomic Nitrogen Use Efficiency (AgNUE), AgNUE values calculated under both high and low nitrogen conditions were plotted, with each point representing a genotype. A color gradient was used to indicate the percentage of nitrogen accumulated in tubers under low nitrogen conditions (%NutLowN). To further explore the relationship between yield and the carbon-to-nitrogen (C/N) balance, total nitrogen and carbon accumulation values under low nitrogen conditions were also plotted, with AgNUE represented through a continuous color scale. These visualizations provide insights into how nitrogen efficiency relates to biomass composition and source– sink dynamics under nitrogen-limited conditions. All plots were generated using the *ggplot2* package in R (Wickham *et al.,* 2009).

### Genotyping and Genome-wide Association Study

We use genotyping by sequencing (GBS) SNP (single nucleotide polymorphism) data obtained by Juyo *et al*. (2019). A new SNP calling was performed based on the latest potato reference genome V6.1 version (Pham *et al.,* 2020), and the resulting SNP matrix is provided in S2 Table. For this, the quality of the reads (generated using the Illumina HiSeq2500 system) was assessed with the FastQC tool (Brown *et al.,* 2017). Subsequently, the reads with a Quality Score (QS) below 30 were cleaned using the Trimmomatic (Bolger *et al.,* 2014), and SNP calling was performed with the NGSEP (Duitama *et al.,* 2014) and TASSEL software (Bradbury *et al.,* 2007). Filtering SNPs with a frequency below 1% and those with missing data less than 15%, resulting in a matrix of 26,515 SNPs. From the VCF file and using the TASSEL software (Bradbury *et al.,* 2007), were filtered those SNPs with minor allele frequency (MAF) ≥ 0.01 and present in at least 85% of the genotypes.

GWAS was performed using the mean values of the 18 phenotypic variables of NUE and the genotyping SNP matrix of 17,374 markers. The associations were conducted using the BLINK (Bayesian Linear Mixed Model) (Huang *et al.,* 2019), MLMM (Mixed Linear Model Method) (Segura *et al.,* 2012) and FarmCPU (Fixed and Random Model Circulating Probability Unification) (Liu *et al.,* 2016) models implemented in the R package ‘Genomic Association and Prediction Integrated Tool’ (GAPIT) (Lipka *et al.,* 2012). Principal components analysis (PCA) for population structure was established and incorporated as correction in the association models. Pairwise correlation coefficients were calculated, and the Linkage Disequilibrium (LD) mean decay was established by plotting the r2 values against the physical distance and visualizing it using TASSEL (Bradbury *et al.,* 2007). The threshold for detecting significantly associated QTN (quantitative trait nucleotide) was set as a P value ≤ 0.05 and correction for false discovery rate (FDR) of multiple test parameter ≤0.1 using the false discovery rate (FDR) B&H procedure (Benjamini *et al.,* 1995) incorporate by GAPIT (Lipka *et al.,* 2012). The identified QTNs were visualized with Manhattan plots and the model adjustments were established by quantile–quantile plots (QQ plots).

### Finding candidate genes associated with nitrogen use efficiency

The genomic positions of the identified QTNs were established at the latest potato reference genome V6.1 version (Pham *et al.,* 2020). Also, whether these regions correspond to intergenic, coding, intronic, or regulatory sequences. For those QTNs located within coding regions, it was determined whether the SNP causes a synonymous or nonsynonymous change in the gene’s codon. Also, the upstream and downstream genomic regions of each QTN were explored, considering the LD decay value in the association panel, to identify candidate genes for NUE in potato. Finally, the functional annotation of the candidate genes was consulted in the SpudBase for potato genome version v6.1, which was generated using data from the Arabidopsis proteome (Lamesch *et al.,* 2012), the PFAM database v32 (El-Gebali *et al.,* 2019), and the Swiss Prot plant protein database (Wu *et al.,* 2006).

## Results

### Significant differences in the phenotypic variables related to Nitrogen Use Efficiency

The genotypes in the association panel showed significant differences in the 18 phenotypic variables related to NUE measured under both high nitrogen and low nitrogen treatments (ANOVA, p <0.01) (S3 Table). The variable SPAD74 values ranged from 13.6 to 48.9 under high nitrogen, with an average of 33.93, while under low nitrogen, values varied between 14.1 and 41.9, with a mean of 27.51, representing an 18.92% decrease (Fig 1A).

**Fig 1.**
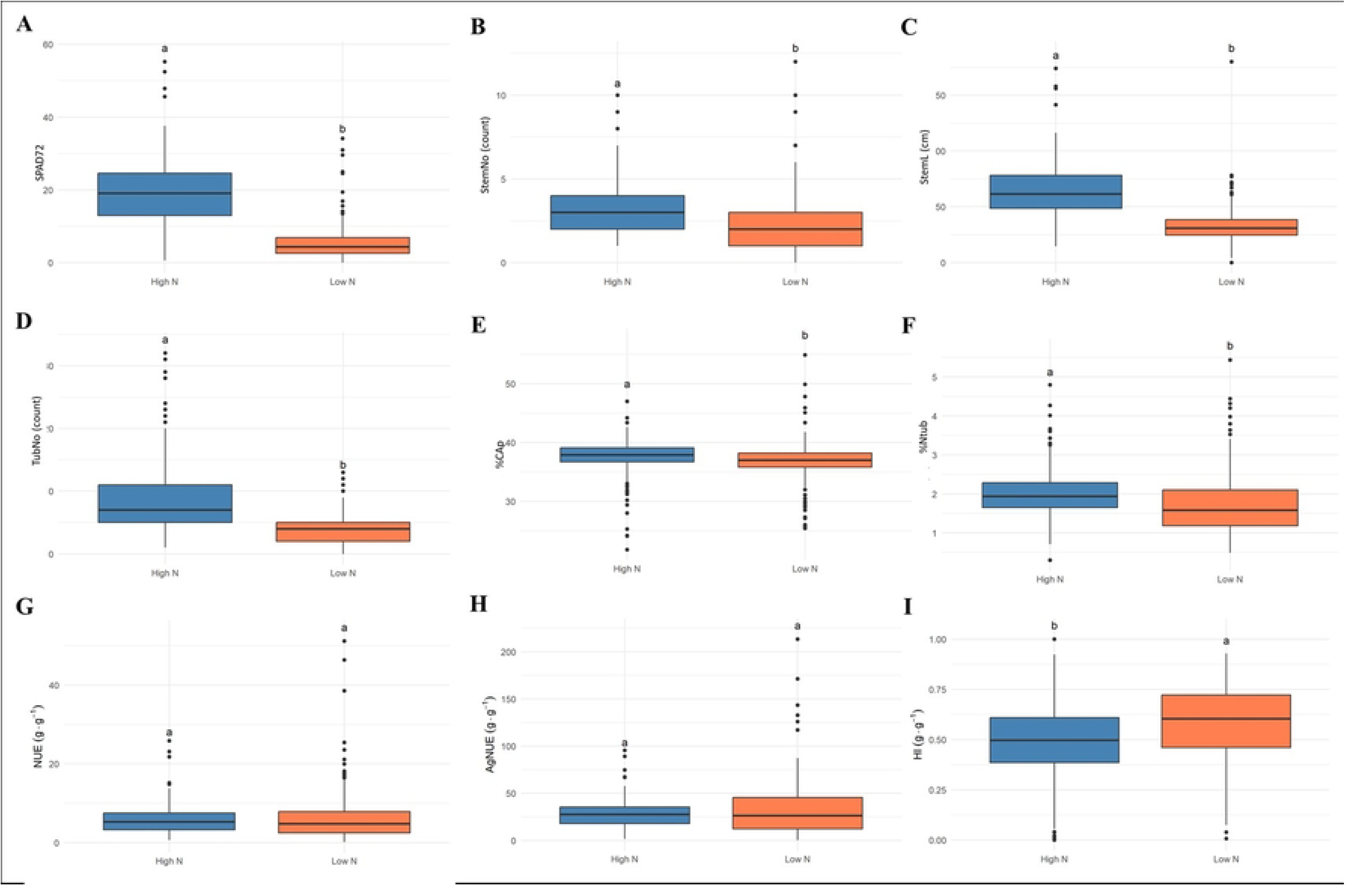
Behavior of phenotypic variables measured for treatments of high and low nitrogen in the potato association panel. A. Chlorophyll Relative Content (SPAD74). B. Number of stems (StemNo). C. Stem length (StemL). D. Number of tubers (TubNo). E. Carbon percentage of the aerial part of the plant (%CAp). F. Tuber N percentage (%Ntub). Nitrogen use efficiency indices (NUE): G. Nitrogen Use Efficiency (NUE, g⋅g^−1^). H. Agronomic Nitrogen Use Efficiency (AgNUE, g⋅g^−1^). I. Harvest Index (HI, g⋅g^−1^). High nitrogen treatment in orange, low nitrogen treatment in blue. The bars indicate the standard deviation (SD), and different letters above each box indicate significant differences according to Tukey’s test (p < 0.05).

Regarding the variable StemNo, this ranged from 1 to 10 under high nitrogen, averaging 2.97, whereas under low nitrogen, values spanned from 1 to 12, with a mean of 2.55, reflecting a 14.14% reduction (Fig 1B). StemL exhibited a broad range, from 14.4 to 174 cm under high nitrogen, with an average of 63.84 cm. Under low nitrogen, values ranged from 4.3 to 180 cm, with a mean of 32.73 cm, indicating a 48.73% decrease (Fig 1C). A similar pattern was observed for TubNo, which varied from 1 to 32 under high nitrogen, averaging 8.57. Under low nitrogen values decreased substantially, ranging from 0 to 13, with a mean of 4.18, corresponding to a 51.22% reduction (Fig 1D).

The organ analysis showed that %CAp under high N ranged from 21.8 to 47%, with an average of 37.63%. Under low nitrogen, values varied between 25.4 and 54.9%, with a mean of 36.9%, showing a 1.94% decline (Fig 1E). The variable %Ntub fluctuated from 0.3 to 4.8% under high nitrogen, with a mean of 2.01%, whereas under low nitrogen, values ranged from 0.49 to 5.44%, averaging 1.72%, resulting in a 14.43% decrease (Fig 1F). The evaluation of NUE indices showed a different trend. NUE ranged from 0.61 to 25.87 g⋅g⁻¹ under high nitrogen, with a mean of 5.8 g⋅g⁻¹. Under low nitrogen, it varied from 0.18 to 51.15 g⋅g⁻¹, with an average of 6.32 g⋅g⁻¹, marking an 8.97% increase (Fig 1G). A similar trend was observed for AgNUE, which ranged from 3.37 to 102.67 g⋅g⁻¹ under high nitrogen, averaging 29.36 g⋅g⁻¹. Under low nitrogen, values were higher, spanning from 0.52 to 213.29 g⋅g⁻¹, with a mean of 32.23 g⋅g⁻¹, indicating a 9.78% increase (Fig 1H). Finally, HI followed an opposite trend, ranging from 0 to 1 g⋅g⁻¹ under high nitrogen, with an average of 0.49 g⋅g⁻¹. Under low nitrogen, values were slightly higher, varying from 0.01 to 0.93 g⋅g⁻¹, with a mean of 0.58 g⋅g⁻¹, reflecting a 19.75% increase (Fig 1I).

The PCA shows that under both low and high nitrogen conditions, the first two components explained 56.6% (Fig 2A: Dim1= 38.7%, Dim2= 17.9%) and 55.6% (Fig 2B: Dim1= 35.2%, Dim2= 20.4%) of the total variation in the phenotypic variables, respectively. The PCA shows that variables such as AgNUE, NUE, HI, TubNo, and StemNo are highly correlated in both treatments. The genotypes are mostly clustered around the center of the plot. Those located on the right-side exhibit high performance such as Col 69, Col 9 and Col 96, with the best performing genotype in terms of yield, Col 93, positioned at the right edge for both treatments. Likewise, the genotype with the lowest yield Col 5 is found on the opposite side of the plot.

**Fig 2.**
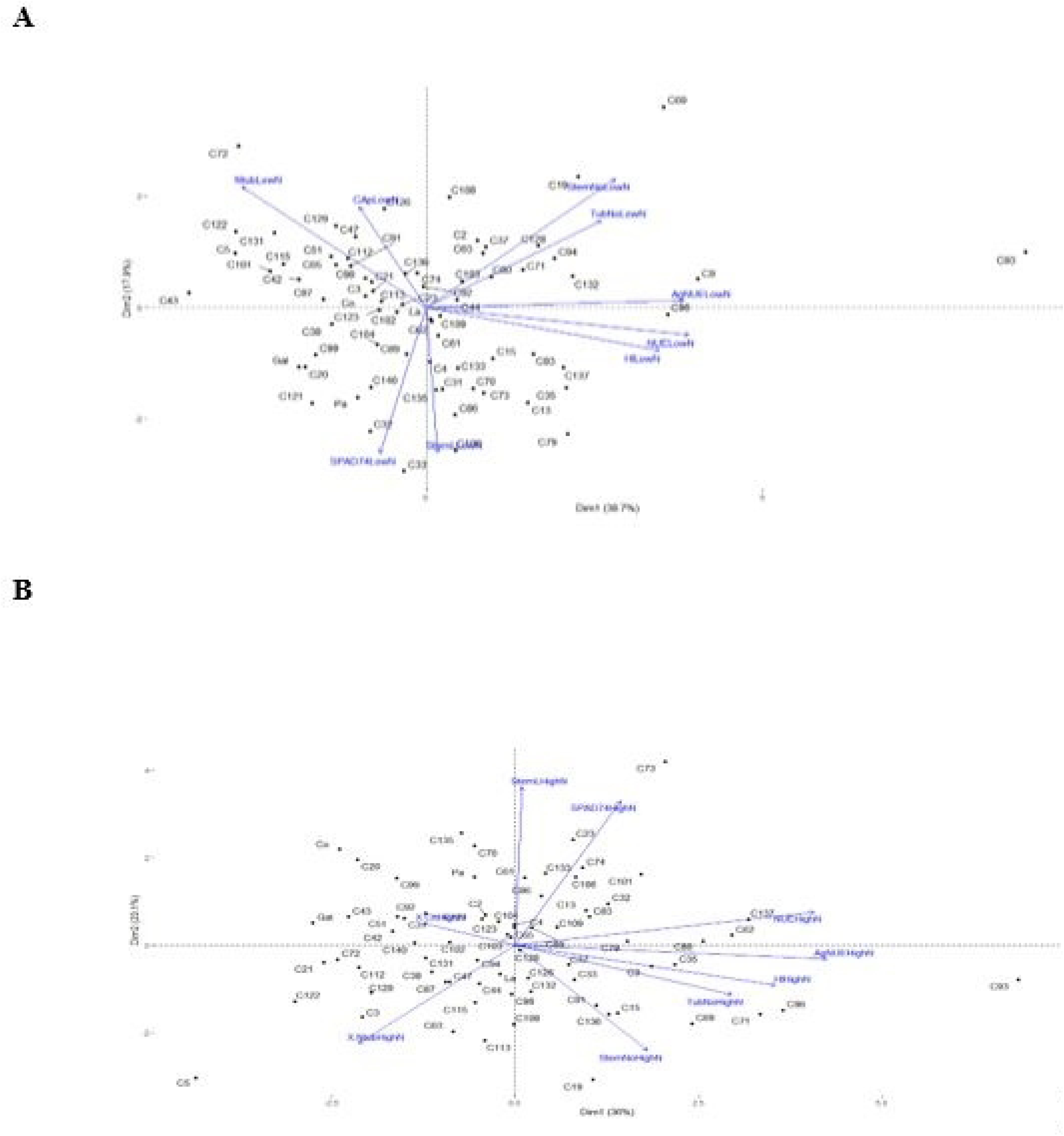
Principal Component Analysis (PCA) of phenotypic variables related to nitrogen use efficiency in the potato association panel. The PCA illustrates the distribution of the phenotypic variables across the first two principal components for **A**. Treatment of low nitrogen. **B**. Treatment of high nitrogen.

In the *ggplot* for AgNUE, genotypes positioned above the regression line represent those with superior performance under nitrogen low conditions (S4 Fig). The genotypes located in the upper right quadrant particularly Col 93, followed by Col 9 and Col 96 demonstrated high AgNUE in both environments, and can be considered efficient under contrasting nitrogen conditions. In contrast, genotypes such as Col 101, Col 115, and Col 91, located in the lower right quadrant, showed good performance under high nitrogen but poor efficiency under low nitrogen. Meanwhile, genotypes like Col 5, Galeras, and Col 21 show a low AgNUE under both nitrogen conditions (S4 Fig). Col 108 has a high percentage of nitrogen in tuber with AgNUE over the percentage of nitrogen in tuber under both conditions.

The results reveal a positive correlation between nitrogen and carbon accumulation in tubers, indicating that both nitrogen and carbon increase proportionally and are strongly associated with AgNUE (S5 Fig). Genotypes with higher nitrogen uptake and utilization accumulate more carbon, due to enhanced photosynthetic capacity and biomass production, where the C/N ratio agrees with the findings of Zheng *et al*. (2018), ranging from 10 to 50. Among all evaluated genotypes, Col 93 shows the highest AgNUE under both nitrogen conditions, nitrogen accumulation, and carbon accumulation, putting it as the most efficient genotype in this study. However, Col 93 also showed a marked decrease in the percentage of nitrogen allocated to tubers, dropping from 1.6% to 0.7% under high and low nitrogen conditions respectively, which negatively impacted tuber nutritional quality.

### Quantitative trait nucleotides (QTNs) identified for nitrogen use efficiency in the association panel

The correction for potential population stratification effects applied to the GWAS models was defined using three principal components (PCs), which together explained 21.05% of the total variance (S6 Fig). Genome Wide Association Study identified a total of 21 QTNs significantly associated with 11 phenotypic variables related to nitrogen use efficiency (Table 1, Fig 3 and S7 Table). QQ plots confirmed a good fit of the association model applied to each phenotypic trait (Fig 4).

**Fig 3.**
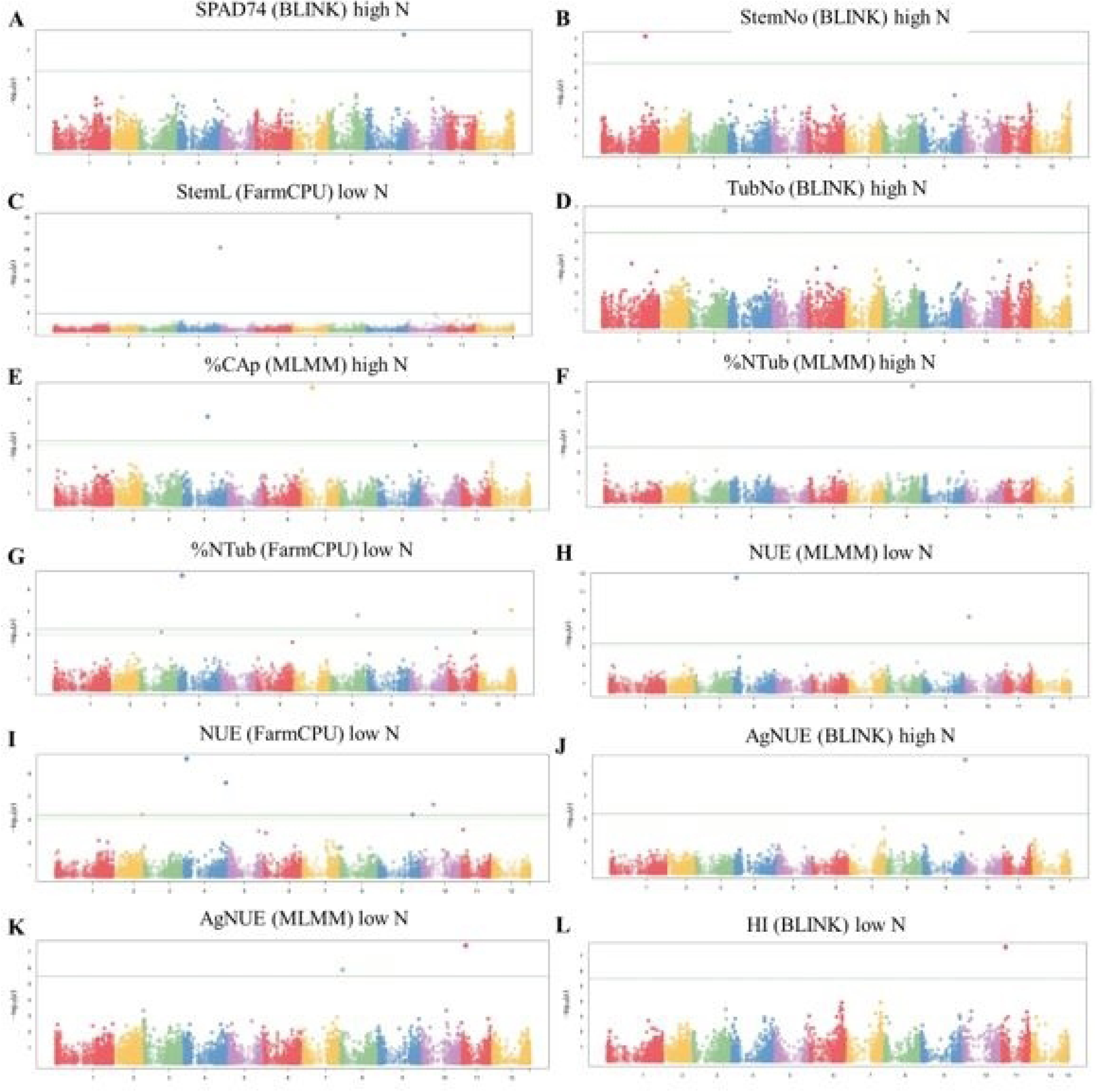
Manhattan plots displaying significant QTNs associated with NUE-related traits in diploid potato under high and low nitrogen conditions, analyzed with different statistical models. **A.** SPAD74 (BLINK) high nitrogen, **B.** StemNo (BLINK) high nitrogen, **C**. StemL (FarmCPU) low nitrogen, **D.** TubNo (BLINK) high nitrogen, **E.** %CAp (MLMM) high nitrogen, **F.** %Ntub (MLMM) high nitrogen, **G.** %Ntub (FarmCPU) low nitrogen, **H.** NUE (MLMM) low nitrogen, **I.** NUE (FarmCPU) low nitrogen, **J.** AgNUE (BLINK) high nitrogen, **K.** AgNUE (MLMM) low nitrogen**, L.** HI (BLINK) low nitrogen. The x axis represents the chromosomal position, while the y axis displays the −log₁₀ (p value) for QTNs associations. The horizontal green line indicates the genome wide significance threshold, with SNPs above this line considered statistically significant.

**Fig 4.**
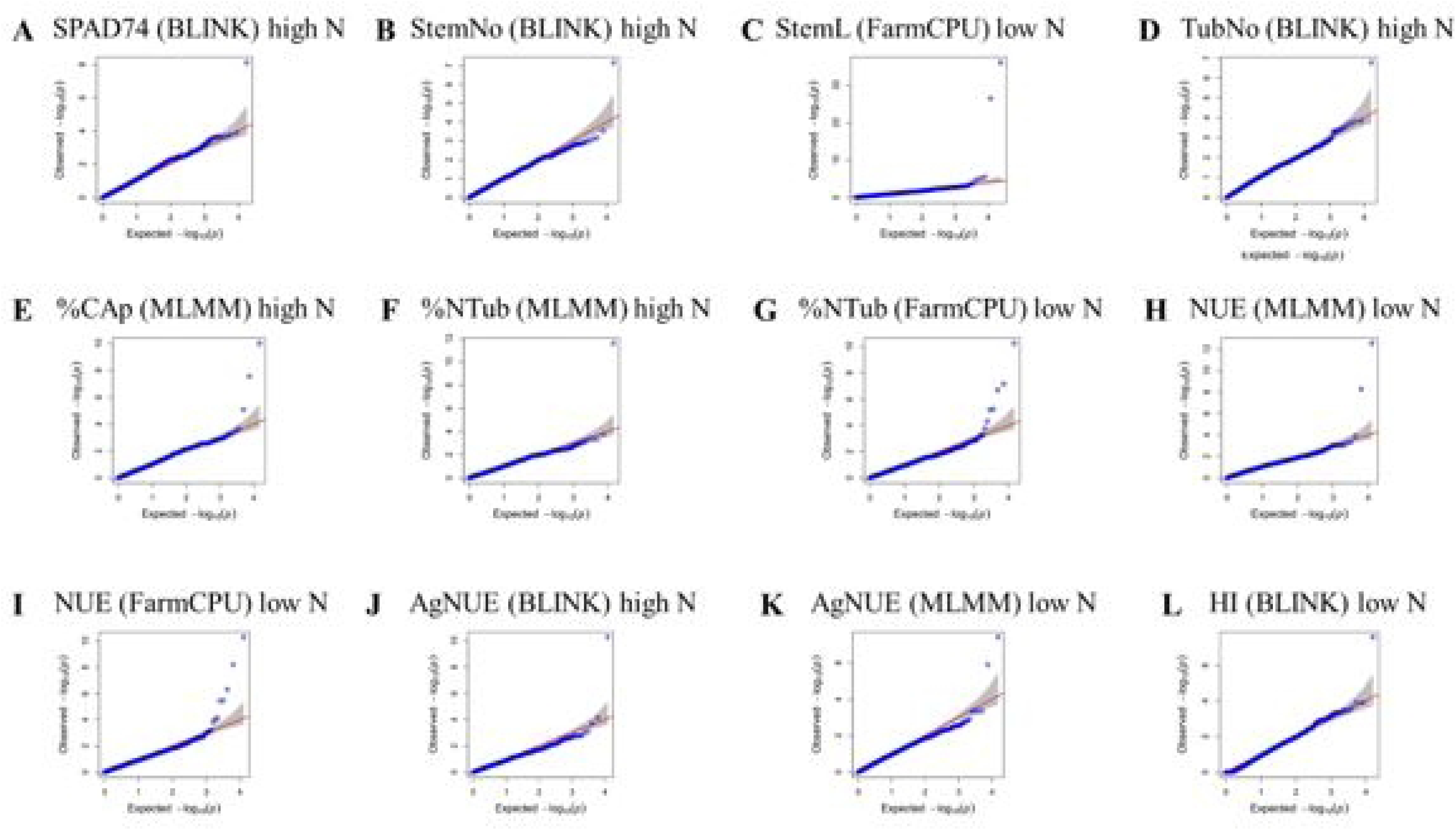
QQ plots showing the goodness of fit of the statistical models used for QTN identification associated with nitrogen use efficiency related traits in diploid potato under high and low nitrogen conditions. **A.** SPAD74 (BLINK) high nitrogen, **B**. StemNo (BLINK) high nitrogen, **C**. StemL (FarmCPU) low nitrogen, **D.** TubNo (BLINK) high nitrogen**, E**. %CAp (MLMM) high nitrogen, **F.** %Ntub (MLMM) high nitrogen, **G.** %Ntub (FarmCPU) low nitrogen, **H.** NUE (MLMM) low nitrogen, **I.** NUE (FarmCPU) low nitrogen, **J.** AgNUE (BLINK) high nitrogen, **K.** AgNUE (MLMM) low nitrogen, **L**. HI (BLINK) low nitrogen. The x axis represents the expected −log₁₀(p value), while the y axis displays the observed −log₁₀(p value) for variant trait associations.

**Table 1.** Summary of QTNs and candidate genes in Linkage Disequilibrium (LD) for phenotypic variables and treatments of nitrogen use efficiency. Phenotypic variables, corresponding treatments of high and low nitrogen (N), total number of identified QTNs, and total number of candidate genes (GC) in linkage disequilibrium (LD) with each QTN.

| Phenotypic variable | Total QTN | p-value range | p-value B&H correction | Total GC in LD |
| --- | --- | --- | --- | --- |
| SPAD74 high N | 1 | $7.50 \times 10^{-09}$ | $1.30 \times 10^{-04}$ | 38 |
| StemNo high N | 1 | $6.70 \times 10^{-08}$ | $1 \times 10^{-03}$ | 40 |
| StemL low N | 2 | $8.8 \times 10^{-37}$ to $3.50 \times 10^{-27}$ | $1.40 \times 10^{-26}$ to $4.20 \times 10^{-19}$ | 74 |
| TubNo high N | 1 | $1.6 \times 10^{-07}$ | $2.6 \times 10^{-03}$ | 53 |
| %CAp high N | 2 | $9.7 \times 10^{-11}$ to $2.80 \times 10^{-08}$ | $1.4 \times 10^{-07}$ to $2 \times 10^{-04}$ | 13 |
| %Ntub high N | 1 | $2.5 \times 10^{-12}$ | $3.6 \times 10^{-08}$ | 31 |
| %Ntub low N | 3 | $5.6 \times 10^{-11}$ to $2.07 \times 10^{-07}$ | $8.2 \times 10^{-07}$ to $1 \times 10^{-03}$ | 107 |
| NUE low N | 6 | $2.8 \times 10^{-13}$ to $3.6 \times 10^{-06}$ | $3.5 \times 10^{-09}$ to $9.23 \times 10^{-03}$ | 202 |
| AgNUE high N | 1 | $5.2 \times 10^{-11}$ | $6.4 \times 10^{-07}$ | 59 |
| AgNUE low N | 2 | $3.8 \times 10^{-08}$ to $1.20 \times 10^{-06}$ | $5.8 \times 10^{-04}$ to $9.50 \times 10^{-03}$ | 88 |
| HI low N | 1 | $2.8 \times 10^{-08}$ | $4.13 \times 10^{-04}$ | 45 |

The variables NUE low nitrogen and %Ntub low nitrogen exhibited the highest number of associated QTNs, with 6 and 3, respectively. The significance of the identified QTNs ranged from p-value of 8.8×10^−37^ to 3.6 x10^−6^, with QTN StemL low nitrogen located at Chromosome 8 position 15,178,651 showing the highest significance and QTN NUE low nitrogen located at Chromosome 8 position 54,180,237, the lowest. The phenotypic variance explained (PVE) by the identified QTNs ranged from 0.08% to 2.73%. The QTN associated with SPAD74 under high nitrogen, exhibited the lowest PVE (0.08%). In contrast, the QTN linked to stem length low nitrogen, showed the highest individual PVE (2.73%). Three additional QTNs related to NUE under low nitrogen conditions explained 1.62%, 1.99%, and 2.29% of the phenotypic variance. These variants together provide a high contribution to the genetic architecture of NUE (Table 1 and S7 Table).

### Repertoire of candidate genes identified for nitrogen use efficiency

From the 21 QTNs identified, nine localize with exonic regions, while 12 localize with intronic regions. Among those located in coding regions, the analysis revealed that all QTNs caused nonsynonymous changes in the gene’s codon, potentially altering protein function (S7 Table). The analysis of LD decay revealed a drop at 263 Kb (S8 Fig). Consequently, this window size around the identified QTNs was used for the search of candidate genes (CGs) in linkage disequilibrium with the associated variable. The total number of CGs identified for the 11 phenotypic variables for which QTNs were detected was 750. The number of CGs ranged from 13 for the %CAp high nitrogen variable to 202 for NUE under low nitrogen conditions (Table 1 and S9 Table).

Within the repertoire of candidate genes, several genes were found whose functional annotation are related to specialized roles in plant nitrogen metabolism. These include genes involved in multiple physiological and regulatory pathways associated with NUE, that can be classified in *nitrogen uptake nitrogen utilization, nitrogen transport and NUE hormonal regulation*. Candidate genes in the *nitrogen uptake category* stand out, such as genes encoding nutrient transporters, including those for nitrogen: (Soltu.DM.12G023520.1, Soltu.DM.05G000760.3), potassium (Soltu.DM.02G032740.1, Soltu.DM.10G001150.1), magnesium (Soltu.DM.01G027980.1, Soltu.DM.05G000840.1), sulfur (Soltu.DM.09G024940.1), zinc (Soltu.DM.10G006310.1), iron (Soltu.DM.09G025190.1), and copper (Soltu.DM.10G000590.1). Also, a *MYB* family member (Soltu.DM.02G033070.1) a gene coding for a phosphate deficiency response (PHR) transcription factor. (S9 Table). Within the *nitrogen utilization* were found CGs coding for an Isocitrate Dehydrogenase (IDH) (Soltu.DM.02G033150.1), an enzyme Ornithine carbamoyltransferase (Soltu.DM.04G035700.1), a gene that encodes a Patatin like protein (Soltu.DM.09G019820.1), and two genes coding for proteins involved in protein degradation (Soltu.DM.11G007670.3) and (Soltu.DM.04G036070.1).

In the *nitrogen transport* category, the gene Soltu.DM.05G000760.3, which encodes a ureide permease, was identified, along with several members of the ATP binding cassette (ABC) transporter family (Soltu.DM.04G007120.1 and Soltu.DM.04G007130.1). Regarding hormonal regulation related to NUE, the candidate gene set included an amidase family protein (Soltu.DM.03G032260.2) and a gene encoding an Indole-3-butyric acid (IBA) response protein (Soltu.DM.12G023410.1). Additionally, genes associated with root hair development and ethylene signaling were identified, such as *Cysteine_synthase_C1* (Soltu.DM.10G006300.1), along with two genes involved in the cytokinin biosynthetic pathway: tRNA isopentenyltransferase (Soltu.DM.10G006560.1 and Soltu.DM.10G006610.1), both encoding isopentenyltransferases. Finally, a gene related to carbon sequestration was identified: Chlorophyll AB binding protein (Soltu.DM.09G025070.1) (Table S4).

## Discussion

Here we are reporting a total number of CGs identified for NUE for the 11 phenotypic variables measured for which QTNs were detected was 750, which contributes to deciphering the genetic structure underlying NUE in *Solanum tuberosum*, one of the world’s most important crops. This is a fundamental step for understanding the genetic control of this trait and a basic contribution to the plant breeding knowledge toward more sustainable and environmentally friendly agriculture. Nitrogen is an essential nutrient for plant growth and productivity and is used in large quantities in agriculture (Hirel & Krapp, 2020). However, its excessive and inefficient use in crops like potato raises environmental concerns due to greenhouse gas emissions, water pollution, and disruption of nearby ecosystems (Getahun *et al.,* 2020; Saravia *et al.,* 2016). Understanding the genetic regulation of NUE and applying this knowledge to potato breeding could reduce fertilizer use, increase production efficiency, and mitigate environmental impacts.

The candidate gene repertoire identified through GWAS in this study reveals a wide array of genomic regions potentially involved in regulating NUE in potato. While future functional validation of these CGs is needed. The current functional annotation of these CGs offers valuable clues to guide such efforts. We focused on four functional categories related to nitrogen metabolism: *nitrogen uptake, nitrogen utilization, nitrogen transport, and hormonal regulation of NUE*.

### Nitrogen uptake

Membrane transporters are key regulators of nitrogen and other essential nutrient uptake and distribution throughout the plant. We identified genes encoding transporters for nitrogen, potassium, magnesium, sulfur, zinc, iron, and cooper. These were associated with nearly all evaluated traits, suggesting NUE in potato depends on coordinated regulation of multiple nutrient transport systems, not only nitrogen specific ones. This aligns with recent evidence on nutrient crosstalk, showing that nitrogen availability can modulate the expression of phosphate deficiency response (PHR) transcription factors members of the MYB family (Fan *et al.,* 2021), such as Soltu.DM.02G033070.1 identified in this study. PHRs regulate phosphorus transporter expression and have been proposed as master regulators integrating nitrogen, phosphorus, sulfur, and iron homeostasis (Kumar *et al.,* 2021), supporting the idea that NUE relies on integrated nutrient uptake regulation. One key CG identified was Soltu.DM.12G023520.1, encoding a nitrate transporter of the NRT1/PTR family. NRT1s are low affinity transporters essential for nitrate uptake and movement across membranes. In rice, overexpression of NRT1 members improves NUE and biomass production (Yang *et al.,* 2023; Fang *et al.,* 2012; Wang *et al.,* 2018). These transporters facilitate cell to cell nitrate movement and enhance soil nitrate uptake, crucial for nitrogen utilization.

### Nitrogen utilization

In the nitrogen assimilation pathway, ammonium is transferred to 2-oxoglutarate to produce glutamate, a key amino acid precursor (Forde *et al.,* 2007). The *IDH* gene (Soltu.DM.02G033150.1), associated with NUE under low nitrogen in our study, is involved in 2-oxoglutarate synthesis (Wei *et al.,* 2023), suggesting it could be a rate limiting enzyme in this pathway. Another gene, Soltu.DM.04G035700.1, encodes ornithine carbamoyltransferase, which uses glutamate to synthesize arginine, a nitrogen storage compound and nitric oxide (NO) precursor, essential for root development (Urbano *et al.,* 2020). We also identified Soltu.DM.09G019820.1, encoding a Patatin like protein that represents over 45% of tuber nitrogen and is known to increase concentration with higher soil nitrogen (Dimante *et al.,* 2024). Additionally, we found genes related to protein degradation, a process critical for nitrogen recycling from leaves. In *Arabidopsis*, overexpression of UBP12 and UBP13 (Soltu.DM.11G007670.3) accelerates senescence under nitrogen deficiency, facilitating nitrogen remobilization to growing tissues (Park *et al.,* 2019). A similar role is played by SAG12 (Soltu.DM.04G036070.1), a papain-like cysteine protease, involved in protein degradation during senescence (Liu *et al.,* 2018).

Col 93 showed a marked decrease in the percentage of nitrogen allocated to tubers, dropping from 1,6% to 0,7% under high and low nitrogen conditions respectively, which negatively impacted tuber nutritional quality. This reduction suggests that most nitrogen corresponds to non-essential storage nitrogen forms, rather than nitrogen required by the growing tuber tissues.

### Nitrogen transport

Allantoin, a ureide derived from nucleic acid degradation, is involved in nitrogen recycling (Soltabayeva *et al.,* 2018). In potato, 10 members of the UPS family have been identified, including the allantoin, they play an important role in nitrogen transport (Huang *et al.,* 2024). In *Arabidopsis*, ureides are transported from senescing to developing tissues by ureide permeases (Lescano *et al.,* 2020). We identified a homologous gene (Soltu.DM.05G000760.3), suggesting a role in long distance nitrogen transport. Proteins are a major nitrogen source in senescing leaves, contributing over 50% to nitrogen remobilization (Soltabayeva *et al.,* 2018). Exported mostly as amino acids, peptides, or small proteins, this transport is facilitated by protein transporters like ATP Binding Cassette (ABC) transporters (Soltu.DM.04G007120.1, Soltu.DM.04G007130.1). These encode H⁺ pumps that drive molecule transport using electrochemical gradients (Rea, 2007). In Arabidopsis, ABC transporters are also crucial for auxin transport, influencing root growth (Jenness *et al.,* 2022). The identification of these transporters as nitrate transporters in cyanobacteria (Li *et al.,* 2024), provides a basis to investigate their potential role in nitrate uptake and transport in potato.

### Hormonal regulation of Nitrogen Use Efficiency

Hormones like auxins and ethylene play key roles in root development, especially under limited nutrient conditions, by improving soil exploration and nutrient uptake. Several CGs involved in hormonal signaling were identified, including an amidase (Soltu.DM.03G032260.2) that catalyzes IAM to IAA an active auxin (Pérez *et al.,* 2021), and an indole-3-butyric acid (IBA) response protein (Soltu.DM.12G023410.1), promoter of root growth in potato (Meenakshi *et al.,* 2021). In *Arabidopsis*, overexpression of AMI1 (amidase family) significantly increases the number of adventitious roots (Pérez *et al.,* 2021; Moya *et al.,* 2021). We also found CGs involved in ethylene signaling and in the regulation of root hair development. Cysteine_synthase_C1 (Soltu.DM.10G006300.1) was associated with NUE under low nitrogen. In *Arabidopsis*, knockout of this gene impairs root development without affecting shoots (García *et al.,* 2010). It also upregulates ethylene related genes like arabinogalactan protein (Soltu.DM.08G004630.1) and ACC synthase (Soltu.DM.08G004500.1). ACC, an ethylene precursor, affects root architecture, especially lateral root elongation (Lee *et al.,* 2019; Yin *et al.,* 2023).

We also found genes in the cytokinin biosynthetic pathway, including tRNA isopentenyltransferase (Soltu.DM.10G006560.1, Soltu.DM.10G006610.1) and a gene cluster (Soltu.DM.10G006390.1 to Soltu.DM.10G006620.1) encoding isopentenyltransferase key enzymes for cytokinin biosynthesis. Cytokinins regulate shoot meristem activity and suppress root meristem activity (Li *et al.,* 2021). In rice, low nitrogen reduces cytokinin levels, promoting seminal root growth via meristem cell proliferation and elongation (Wang *et al.,* 2020). Multiple gene copies suggest a duplication event that could enhance cytokinin production and influence root development under nitrogen limited conditions.

In this study, ammonium nitrate was used as the nitrogen source, beyond its nutritional role, nitrate is also recognized as a signaling molecule that induces cytokinin biosynthesis in roots, as extensively reported in the literature (Sakakibara *et al.,* 2003) and the presence of multiple related genes may reflect a enhances cytokinin production, potentially influencing root development.

A gene associated with carbon sequestration in tubers, and NUE was also identified: the chlorophyll AB binding protein (Soltu.DM.09G025070.1), linked to photosystems I and II. This protein captures and transfers excitation energy under different light intensity and helps balance energy distribution, increasing light use efficiency and potentially enhancing carbon assimilation by leaves (Sultana *et al.,* 2020; Zhang *et al.,* 2024). This accumulation of carbon could give an advantage to Col 93 in terms of dry matter accumulation in tubers compared to the other genotypes.

Given NUE has strong environmental dependency, accurate phenotyping is essential. We applied a substrate grow system under low nitrogen conditions to precisely assess NUE related traits at key growth stages. This method can serve as a reference for future research in crops with similar agronomic traits. Incorporating high throughput phenotyping, especially in the field, would further refine physiological measurements. Finally, the QTNs and CGs identified through GWAS offer promising targets for potato breeding. Prioritization and validation of top CGs via gene editing or overexpression could clarify their roles in NUE and support the development of potato varieties with lower fertilizer needs and contribute to sustainable agriculture.

The genotypes Col 5, Galeras, and Col 21 exhibited low nitrogen use efficiency under both high and low nitrogen conditions. One possible explanation is that these accessions were not selected for yield related traits but rather preserved due to their genetic diversity. Consequently, they do not appear to carry favorable alleles associated with efficient nitrogen assimilation and remobilization. In our GWAS analysis, we identified several candidate genes involved in nitrate transport (NRT family), carbon-nitrogen transport, and root development, which showed weak or no association with these genotypes. This could partially explain their low performance under both nitrogen regimes.

## Conclusion

The QTNs and candidate genes identified represent a major step forward in further dissecting the NUE in potato. Nevertheless, the polygenic nature of the trait implies that it is influenced by the interaction of multiple genes whose roles are determined by different physiological mechanisms. Thus, the findings described here require further research and functional validation efforts to contribute to our understanding of the underlying biological process but also to allow, in the near future, the development of cultivars with optimized nitrogen yield, reducing the need for synthetic fertilizers, which is especially important in the global context of rising food demand and restrictions on the use of agricultural inputs.

## Acknowledgments

The authors thank the Sistema General de Regalías of Colombia and Minciencias for funding the project entitled “Aprovechamiento de la biodiversidad en agraz y papa para el desarrollo de cultivos promisorios en el departamento de Santander”, identified with No. BPIN 2020000100075 and belongs the call “Convocatoria para la conformación de un listado de propuestas de proyectos elegibles para el fortalecimiento de capacidades institucionales y de investigación de las instituciones de educación superior públicas”.

## Author contributions

**Conceptualization:** TMV, JCSS, MCDN.

**Data curation:** MAMB, ANJM, JCSS.

**Formal analysis:** MAMB, ANJM, JCSS, GALM, SM.

**Project administration:** TMV, JCSS, MCDN

**Writing – original draft:** MAMB, ANJM, JCSS.

**Writing – review & editing**: All authors

## Supporting information

**S1 Table**. Association panel

**S2 Table**. SNPs matrix.

**S3 Table.** ANOVA of the variables evaluated under high and low N conditions.

**S4 Fig.** Relationship between Agronomic Nitrogen Use Efficiency (AgNUE) under high and low nitrogen conditions. Each point represents a genotype. The color gradient indicates the percentage of nitrogen accumulated in tubers under low nitrogen conditions (%NtubLowN), with green tones indicating lower values and darker tones higher values.

**S5 Fig.** Relationship between accumulated carbon and nitrogen in tubers under low nitrogen conditions. Each point represents a genotype. The color gradient indicates Agronomic Nitrogen Use Efficiency (AgNUE), with green tones representing lower values and darker tones higher values.

**S6 Fig.** PCA for population structure.

**S7 Table.** List of QTNs identified by GWAS models.

**S8 Fig.** Linkage disequilibrium (LD) decay.

**S9 Table.** Repertoire of candidate genes.

## Data Availability Statement

All supporting data for this research are provided in the Supplementary Materials.

